# Glucose uptake in pigment glia suppresses tau-induced inflammation and photoreceptor degeneration in *Drosophila*

**DOI:** 10.1101/2024.08.14.607919

**Authors:** Mikiko Oka, Sho Nakajima, Emiko Suzuki, Shinya Yamamoto, Kanae Ando

## Abstract

Brain inflammation contributes to the pathogenesis of neurodegenerative diseases such as Alzheimer’s disease (AD). Glucose hypometabolism and glial activation are pathological features seen in AD brains; however, the connection between the two is not fully understood. Using a *Drosophila* model of AD, we identified that glucose metabolism in glia plays a critical role in neuroinflammation under disease conditions. Expression of human Tau in the retinal cells, including photoreceptor neurons and pigment glia, causes photoreceptor degeneration accompanied by inclusion formation and swelling of the lamina cortex. We found that inclusions are formed by glial phagocytosis, and swelling of the laminal cortex correlates with the expression of antimicrobial peptides. Co-expression of human glucose transporter 3 (*GLUT3*) with Tau in the retina does not affect tau levels but suppresses these inflammatory responses and photoreceptor degeneration. We also found that expression of *GLUT3*, specifically in the pigment glia, is sufficient to suppress inflammatory phenotypes and mitigate photoreceptor degeneration in the tau-expressing retina. Our results suggest that glial glucose metabolism contributes to inflammatory responses and neurodegeneration in tauopathy.

**Summary Statement:** Glucose uptake into pigment glia suppresses inflammatory responses and photoreceptor degeneration in the fly model of tauopathy.

## Introduction

Alzheimer’s disease (AD) is a progressive neurodegenerative disorder and the most common cause of dementia among older populations (Knopman et al., 2021). AD is characterized by the extracellular deposition of β-amyloid and intracellular accumulation of abnormally phosphorylated forms of the microtubule-associated protein tau (Samudra et al., 2023). Neuroinflammation is another core pathology in AD and other neurodegenerative disorders (Sobue et al., 2023). Inflammatory responses in the central nervous system are mediated by mainly glial cells, including microglia and astrocytes, to protect against infections and injuries; however, under neurodegenerative disease conditions, glial cells lose their neuroprotective functions and acquire reactive phenotypes (Ransohoff, 2016; Streit, 2002). Although they can remove protein aggregates by phagocytosis, activated glial cells can also harm neurons via excess phagocytosis and the release of neurotoxic cytokines (Chen and Holtzman, 2022). However, what triggers glial activation under disease conditions is not fully understood.

A decline in brain glucose metabolism has been pathologically linked to AD (Butterfield and Halliwell, 2019). Most of the ATP required to support brain function is supplied by glucose metabolism (Bélanger et al., 2011), and glucose utilization is reduced in the brains of AD patients (Dukart et al., 2013; Huang et al., 2020; Oka et al., 2021). Dysregulated glucose metabolism in AD may be due to cerebral hypoperfusion (Park et al., 2019) or downregulation of glucose transporters (GLUTs) that mediate glucose uptake across the blood-brain barrier and delivery to glial cells (Bélanger et al., 2011; Koepsell, 2020; Kyrtata et al., 2021; Liu et al., 2008). Glial cells also rely on glucose, and cellular metabolism can regulate their activation (Wang et al., 2019; Xiang et al., 2021). The mitochondrial oxidative phosphorylation is reduced in the activated microglial cells (Nair et al., 2019), and mitochondrial dysfunction in astrocytes has been reported in amyotrophic lateral sclerosis and neuroinflammation models (Fiebig et al., 2019; Martínez-Palma et al., 2019). While these reports suggest that targeting glial metabolism may be a strategy to decrease neuroinflammation and suppress neurodegeneration, mechanistic investigation in an *in vivo* model is required to explore this possibility.

Here, we investigated the roles of glucose metabolism in glial cells on tau-induced neurodegeneration in a *Drosophila* model. We identified glia-associated phenotypes in the fly retina expressing human Tau, such as inclusion-like structures by phagocytosis and induction of antimicrobial peptide (AMP) expression. *Drosophila* retina contains neurons and several glial cells, and we found that pigment glia plays a critical role in tau-induced photoreceptor degeneration. Enhancement of glucose uptake in pigment glial cells ameliorates tau-induced laminal cortex swelling as well as photoreceptor degeneration. Our results suggest that glucose hypometabolism in glial cells contributes to neurodegeneration in tauopathy.

## Results

### Tau expression in the fly retina causes photoreceptor degeneration, swelling of the laminal cortex, and inclusion formation

Expression of human Tau in *Drosophila* using the pan-retinal *GMR-GAL4* driver causes a rough eye phenotype in the compound eye surface accompanied by the formation of vacuoles caused by apoptosis in the retina and axonal degeneration in the lamina (Fig. 1A) (Iijima-Ando et al., 2012; Jackson et al., 2002; Wittmann et al., 2001). UAS-tau transgene alone does not cause these phenotypes as described previously (Wittmann et al., 2001 and Fig. S1). Transmission electron microscopy (TEM) revealed degenerating photoreceptor neurons with disoriented rhabdomere in tau-expressing flies (Fig. 1B and C, an ommatidium colored in yellow). We also noticed swelling of the laminal cortex (shaded in blue in Fig. 1A), which is located between the lamina and retina and consists of layers of L1-L5 neurons, surface glia (perineurial glia (PNG) and subperineurial glia (SPG)) and cortex glia (Chotard and Salecker, 2007; Kremer et al., 2017). We analyzed the neuronal and glial cell distribution in the laminal cortex by using glial expression of mCherry carrying the nuclear localization signal (mCh::NLS) and immunostaining with neuronal marker, elav (Fig. 1D). In the laminal cortex, surface glia and cortex glia form two layers, and cell bodies of L1-L5 neurons form another layer. With tau expression in the retina with GMR promoter, the neuronal layers in the laminal cortex were enlarged with an increased number of neurons. In contrast, glial cells in the laminal cortex were degenerated (Fig. 1D).

**Figure 1.**
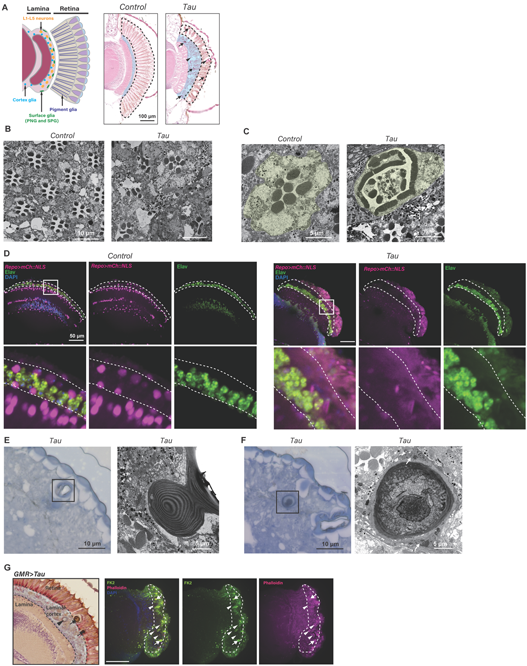
Human tau expression in the fly retina causes photoreceptor degeneration, swelling of the laminal cortex, and inclusion formation. (A) Schematic representation of the retina (Left) and hematoxylin and eosin (H&E)-stained sections (Right). Human tau expression causes degeneration, which is indicated by vacuoles in the retina (dashed line), swelling of the laminal cortex (blue shade), and inclusions (arrows and arrowheads). Tau expression was driven by *GMR-GAL4*, and luciferase was used as a control (Control). The flies were 5 days old. Scale bar, 100 µm. (B-C) Representative transmission electron microscopy (TEM) images of eyes in 5-day-old flies expressing luciferase (*Control*) or human Tau (*Tau*). Tau expression showed photoreceptor degeneration with disoriented rhabdomere (yellow shade). (D) Tau expression increases neurons and reduces glial cells in the laminal cortex. Glial cells were labeled with Repo>mCh::NLS (magenta), and neurons were immunolabeled with an anti-elav antibody (green). DAPI was used to stain nuclei (blue). To analyze the effect of tau expression in the retinal cells, *GMR-tau (P301L),* which expresses human Tau with pathological mutation (P301L) directly under the control of the GMR regulatory sequence, was combined with Repo>mCh::NLS (Tau). The laminal cortex is outlined by dotted lines. Bottom: blow-up images. Scale bar: 50 µm. (E-F) Toluidine blue staining and TEM images of inclusion in the Tau-expressing retina. Tau expression showed two types of inclusions: invaginated corneal lens (E, arrowheads in A) and debris-containing inclusions wrapped by glial cells (F, arrows in A). (G) (Left) A paraffin-embedded section of a fly retina. The arrowhead points to the invaginated corneal lens, while the arrows indicate the densely dark inclusions. (Right) Immunostaining of Tau-expressing retina with anti-ubiquitin antibody (FK2, green, bottom). Filamentous actin was stained with phalloidin (magenta), and nuclei were stained with DAPI (blue). The laminal cortex was indicated by the dotted line. Scale bar, 50 µm

Tau expression caused the formation of dense inclusion-like structures in the retina and lamina (Fig. 1A). TEM analysis of these structures revealed two distinct ultrastructures: invaginated corneal lens (arrowheads in Fig. 1A, E, and G) and dark inclusions containing cellular debris (arrows in Fig. 1A, F and G) and wrapped by glia (white arrows in Fig. 1F). These inclusions contain ubiquitinated proteins and cytoskeleton such as filamentous actin (Fig. 1G), suggesting that they are dead cells with abnormal protein accumulation and engulfed by glia. These results suggest that glial cells are also affected by tau expression.

### Tau expression induces glial phagocytosis and AMP expression

The appearance of inclusion-like structures (Fig. 1F) was similar to the glial bodies (Dutta et al., 2016), which are membranous structures representing glial hyper-wrapping. We were motivated to test if these inclusions are formed via glial phagocytosis. The glial phagocytic activities in the adult fly brain are mediated by two glial transmembrane phagocytic receptors, Draper (Drpr) and Nimrod C4 (NimC4, also known as SIMU) (Kurant et al., 2008; Scheib et al., 2012). We found that blocking phagocytosis in the retina by RNAi-mediated knockdown of *drpr* and *NimC4* did not affect the eye size or photoreceptor degeneration significantly but reduced the number of inclusions in the tau-expressing retina (Fig. S2 and Fig. 2A). These results indicate that the inclusions are formed by glial engulfment activity.

**Figure 2.**
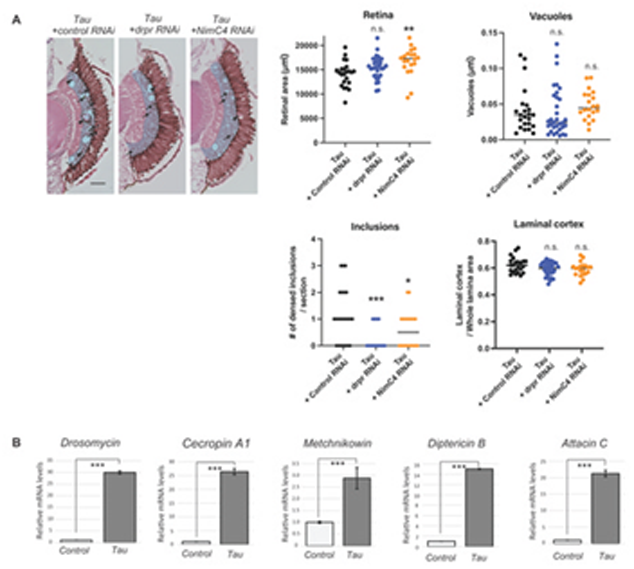
Tau expression induces glial phagocytosis and AMP expression. (A) Inclusions in the tau-expressing retina are formed by glial phagocytosis. RNAi-mediated knockdown of *drpr* or *NimC4* reduced the number of inclusions in the tau-expressing eyes. Representative images of H&E-stained tissue sections and quantification. Scale bar, 100 µm. Mean±SE, n=20-34, ***; p<0.001, **; p<0.01, *; p<0.05, n.s.; p>0.05, One-way ANOVA followed by Tuckey’s HSD multiple comparisons test. Flies were 7 days after eclosion. (B) Tau induces antimicrobial peptide (AMP) expression. Heads carrying *GMR-GAL4* driven luciferase (Control) or human Tau (Tau) were subjected to qRT-PCR to analyze the levels of AMPs. Mean±SE, n=3, ***; p<0.001; Statistical significance was assessed with an unpaired two-tailed t-test. Flies were 7 days after eclosion.

Proinflammatory responses in *Drosophila* involve the expression of antimicrobial peptides (AMPs) downstream of Toll and Immune deficiency (Imd) pathways (Harnish et al., 2021; Hedengren et al., 1999; Sakakibara et al., 2023; Shukla et al., 2019). We found that *GMR-GAL4* driven tau-expressing flies exhibited significantly elevated expression levels of AMPs, including *Drosomycin (Drs)*, *Cecropin A1 (CecA1), Metchnikowin (Mtk), Diptericin B (DptB), and Attacin C (AttC)* (Fig. 2B). These results suggest that tau expression in the fly retina induces glial phagocytosis and inflammatory responses, which are hallmarks of glial activation.

### Enhancement of glucose uptake suppresses tau-induced photoreceptor degeneration and swelling of the laminal cortex

We analyzed the effects of enhancement of glucose uptake in tau-induced phenotypes in the retina by expressing human GLUT3 encoded by the *SLC2A3* gene, which has been shown to enhance glucose uptake in cells effectively (Besson et al., 2015; Manzo et al., 2019; Oka et al., 2021). Co-expression of GLUT3 increased the eye size in tau flies (Fig. 3A, compare *Tau and Tau+GLUT3*, p<0.001). The eyes co-expressing Tau and mCD8::ChRFP were similar to those in flies expressing Tau alone, indicating that the increase in the eye size caused by co-expression of *GLUT3* is not due to non-specific effects of an exogenous protein or dilution of GAL4 activity (Fig. 3A, compare *Tau* and *Tau+mCD8::ChRFP*, p>0.05). The expression of GLUT3 alone did not affect eye size (Fig. 3A, compared to Control and *GLUT3*, p>0.05). Rhodopsin 1 (Rh1) protein level is significantly lower in Tau flies, likely due to loss of photoreceptor neurons (Fig 3B, compare control and *Tau*). Co-expression of GLUT3 restored Rh1 protein levels (Fig. 3B, compare *Tau* and *Tau+GLUT3*, p<0.001). The vacuoles in the retina of flies co-expressing Tau and GLUT3 were also significantly less than those expressing Tau alone (Fig. 3C-D, p<0.01). At ultrastructural levels, while tau-expressing retina contains degenerating photoreceptor neurons with disoriented rhabdomere in tau-expressing flies (Fig. 1C, an ommatidium colored in yellow), ommatidium from flies co-expressing Tau and GLUT3 retained rhabdomeres with proper orientation (Fig. 3F, *Tau+GLUT3*). Co-expression of GLUT3 did not affect the number of inclusion-like structures (Fig. 3E), while suppressed laminal cortex swelling (Fig. 3F). Overexpression of *Drosophila* Glut1 (Fig. S3), a fly homolog of human GLUT3, also suppressed tau-induced retinal degeneration and the lamina cortex swelling (Fig. 3G-I). These results indicate that glucose uptake enhancement suppresses tau-induced neurodegeneration and glial phenotypes.

**Figure 3.**
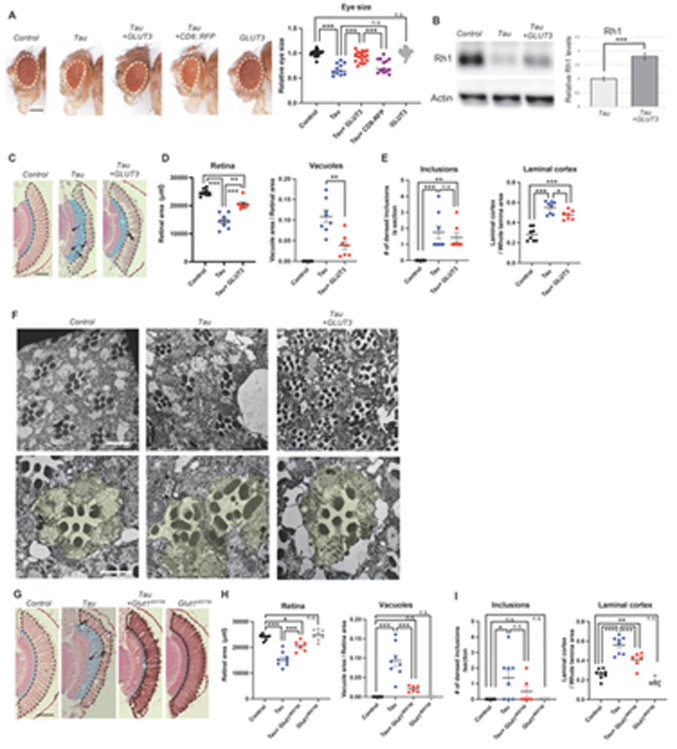
Enhancement of glucose uptake suppresses tau-induced photoreceptor degeneration and swelling of the laminal cortex. (A) The reduction in the eye size caused by tau expression (compare Control and *Tau*) was suppressed by GLUT3 expression (compare *Tau* and *Tau+GLUT3)*. Expression of a control protein mCD8::ChRFP did not affect tau-induced eye phenotype (compare *Tau* and *Tau+mCD8::ChRFP*), and expression of GLUT3 alone did not affect eye size (compare *Control and GLUT3*). Scale bar, 250 µm. Mean±SE, n=11-27, ***; p<0.001, n.s.; p>0.05, One-way ANOVA followed by Tuckey’s HSD multiple comparisons test. (B-F) Human GLUT3 expression suppresses tau-induced photoreceptor degeneration and glial phenotype. (B) GLUT3 expression suppressed tau-induced reduction in Rhodopsin 1 protein levels (Rh1). Fly heads expressing Tau with a control protein mCD8::ChRFP (*Tau*) or Tau and GLUT3 (*Tau + GLUT3*) were subjected to western blotting with an anti-Rh1 antibody. Actin was used as a loading control. Representative blot and quantification are shown. Mean±SE, n=3, n.s., ***; p<0.001, Statistical significance was assessed with unpaired two-tailed t-test. (C-F) GLUT3 co-expression suppressed tau-induced neurodegeneration and glial phenotypes. (C) Representative images of sections. (D) GLUT3 co-expression suppressed tau-induced retinal degeneration. Quantitation of areas of the retina vacuole (Mean±SE, n=7-9, **; p<0.01, ***; p<0.001, Statistical significance was assessed with unpaired two-tailed t-test). (E) GLUT3 co-expression did not affect inclusions but suppressed swelling of the laminal cortex (Mean±SE, n=7-8, *; p<0.05, n.s.; p>0.05, Statistical significance was assessed with unpaired two-tailed t-test) (F) Representative TEM images of photoreceptors. GLUT3 co-expression mitigated rhabdomere disorientation in tau-expressing eyes (yellow shade). (G-I) *Drosophila Glut1* expression also suppressed tau-induced retinal degeneration and swelling of the laminal cortex. Representative images of retinal sections (G) and quantification of areas of the retina and vacuole (H), the number of inclusions, and size of the laminal cortex (I). Mean±SE, n=6-8, *; p<0.05, **; p<0.01, ***; p<0.001, One-way ANOVA followed by Tuckey’s HSD multiple comparisons test (G and I). Flies were 7 days after eclosion. Scale bar, 100 µm.

### Enhancement of glucose uptake does not affect tau phosphorylation and accumulation

To understand the mechanisms underlying the neuroprotective effects of enhanced glucose uptake, we first analyzed the effect of GLUT3 expression on tau phosphorylation and accumulation by western blot with monoclonal antibodies. Co-expression of GLUT3 did not affect the levels of total Tau (Total Tau) or its phosphorylation at disease-associated sites, including Ser262 (pSer262), Ser396/404 (PHF1), and Ser202 (CP13) (Fig. 4). These results suggest that enhanced glucose uptake mitigates retinal degeneration downstream of tau phosphorylation and accumulation.

**Figure 4.**
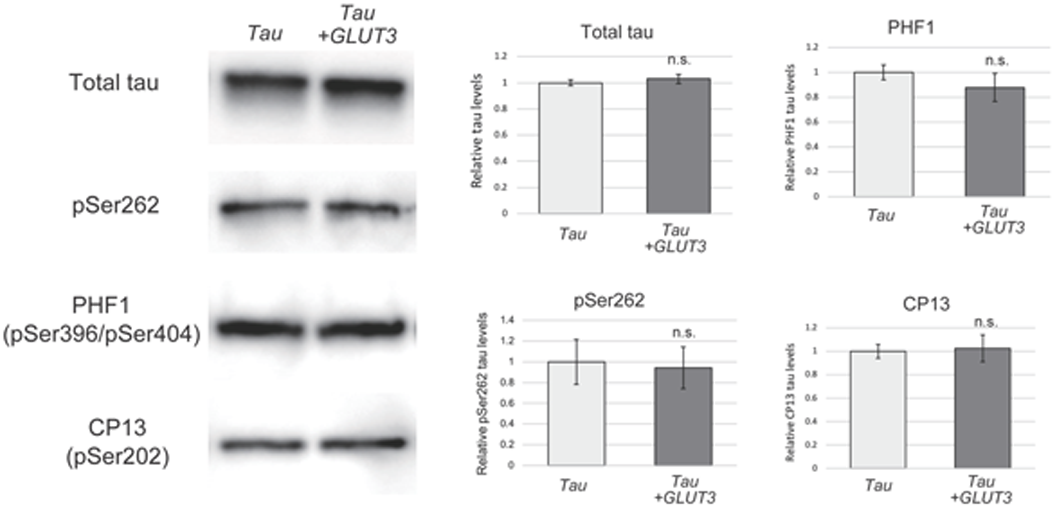
Enhancement of glucose uptake does not affect tau phosphorylation and accumulation. Fly heads expressing human Tau with *CD8::GFP* as a control (Tau) or those co-expressing tau and *GLUT3* (*Tau + GLUT3*) were subjected to western blotting with anti-tau (total Tau) or anti-phospho-tau monoclonal antibodies (pS262, PHF1 and CP13). Actin was used as a loading control. Mean±SE, n=4-6, n.s., Statistical significance was assessed with unpaired two-tailed t-test. Flies were 3 days after eclosion.

### Enhancement of glucose uptake in pigment glial cells suppresses tau-induced neurodegeneration

GMR element used in *GMR-GAL4* drives expression in all the cells behind the morphogenetic furrow, including pigment glial cells (Freeman, 1996). Pigment glial cells surround photoreceptor neurons and support their functions by providing nutrition and uptaking toxic materials (Liu et al., 2017). Expression of *GMR-GAL4* and *54C-GAL4* in pigment glial cells was confirmed by using *UAS-mCh::NLS* (Fig. S4). Although tau expression in the pigment cells does not cause a rough-eye phenotype (Fig. S5), GLUT3 expression in the pigment cells may mediate the protective effects against photoreceptor degeneration. To dissect the effects of GLUT3 expression in glial cells in tau-induced retinal degeneration, we used *GMR-tau (P301L),* which expresses human Tau carrying pathological mutation (P301L) directly under the control of the GMR regulatory sequence (Jackson et al., 2002), in combination with *GAL4/UAS* system to express *GLUT3* using *54C-GAL4*, a pigment glia-specific driver (Nagaraj and Banerjee, 2007). We found that swelling of the laminal cortex (Fig. 5A) and AMP expression (Fig. 5B) was suppressed by the expression of *GLUT3* in pigment glia. Intriguingly, pigment cell-expression of *GLUT3* in the tau-expressing retina also suppressed photoreceptor degeneration (Fig. 5A and C). Vacuole formation was suppressed, eye size was significantly increased, and Rh1 protein level was restored upon *GLUT3* expression in the pigment glia (Fig. 5A, C, and D). In contrast, expression of *GLUT3* in photoreceptor neurons by *Rh1-GAL4* did not mitigate photoreceptor degeneration (Fig. 5E-G), indicating that the protective effects of *GLUT3* co-expression against tau toxicity observed with *GMR-GAL4* (Fig 3) is mediated by pigment glia. These results suggest that enhancement of glucose uptake in pigment glia mitigates inflammatory responses and tau-induced photoreceptor degeneration.

**Figure 5.**
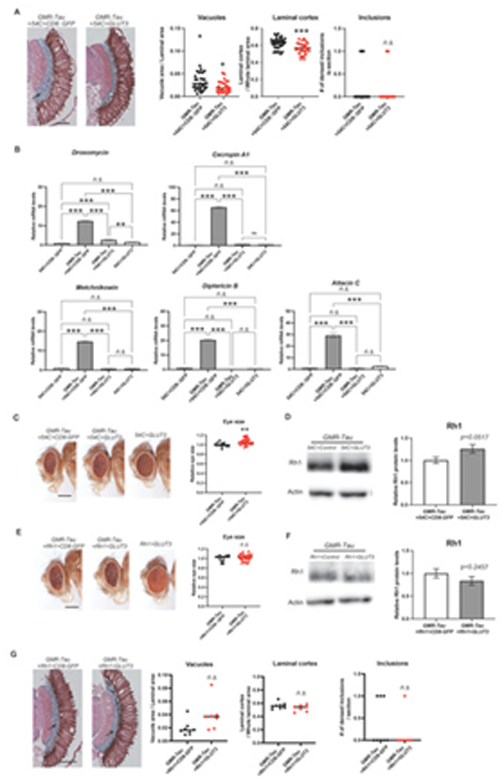
Enhancement of glucose uptake in the pigment glia suppresses tau-induced neurodegeneration. (A-D) Pigment glia-specific GLUT3 expression using *54C-GAL4* rescued neurodegeneration and glial phenotypes in GMR-tau flies. GMR-tau, which expresses human Tau directly under the control of the GMR regulatory sequence, was combined with GLUT3 expression driven by *54C-GAL4*, a pigment glia-specific driver. CD8::GFP was used as a control for GLUT3. (A) Pigment glia-specific GLUT3 expression suppressed photoreceptor degeneration (Vacuoles) and laminal cortex swelling (Laminal cortex) (Mean±SE, n=29-34, *; p<0.05, **; p<0.01, Statistical significance was assessed with unpaired two-tailed t-test, Scale bar, 100 µm). Flies were 7 days after eclosion. (B) GLUT3 expression in the pigment glia suppressed AMP expressions in GMR-tau flies. AMP expression levels were measured by qRT-PCR (B, Mean±SE, n=3, **; p<0.01, ***; p<0.001, One-way ANOVA followed by Tuckey’s HSD multiple comparisons test). (C-D) GLUT3 expression in the pigment cells suppressed the loss of photoreceptor neurons in GMR-tau flies. (C) Sizes of external eyes (Mean±SE, n=12-36, *; p<0.05, Statistical significance was assessed with unpaired two-tailed t-test, Scale bar, 250 µm) and (D) Rh1 protein levels analyzed by Western blotting (Mean±SE, n=3, p=0.0517, Statistical significance was assessed with unpaired two-tailed t-test). (E-G) Photoreceptor neuron-specific GLUT3 expression using *Rh1-GAL4* did not suppress photoreceptor degeneration in GMR-tau flies. GMR-tau, which expresses human Tau directly under the control of the GMR regulatory sequence, was combined with GLUT3 expression driven by *Rh1-GAL4*, a photoreceptor neuron-specific driver. CD8::GFP was used as a control for GLUT3. (E) Sizes of external eyes (Mean±SE, n=13-30, n.s.; p>0.05, Statistical significance was assessed with unpaired two-tailed t-test, Scale bar, 250 µm), (F) Rh1 protein levels analyzed by Western blotting (Mean±SE, n=3, p=0.2457, Statistical significance was assessed with unpaired two-tailed t-test) (G) Photoreceptor neuron-specific GLUT3 expression did not suppress photoreceptor degeneration (Vacuoles) and glial swelling (Laminal cortex) (Mean±SE, n=7-8, n.s.; p>0.05, One-way ANOVA followed by Tuckey’s HSD multiple comparisons test, Scale bar, 100 µm).

## Discussion

Enhancing glucose uptake by neuronal overexpression of GLUTs has been reported to ameliorate neurodegeneration in several neurodegenerative conditions such as huntingtin, β-amyloid, and TDP-43 toxicity in *Drosophila* (Besson et al., 2015; Manzo et al., 2019; Niccoli et al., 2016; Cabirol-Pol et al., 2018) as well as in age-related neuronal dysfunction (Oka et al., 2021). However, the beneficial effect of enhancing glucose uptake in glial cells has not been examined. In this study, we characterized glial phenotypes in the fly retina expressing human Tau and examined the effects of enhanced glucose uptake in the pigment glia on them. We found that tau expression causes phenotypes mediated by glia, such as inclusion formation, swelling in the laminal cortex, and AMP expression (Fig. 1-3). We also found that the expression of glucose transporters in the pigment glia suppresses many of these phenotypes and mitigates photoreceptor degeneration downstream of Tau (Fig. 4 and 5). These results suggest that glucose metabolism in glial cells contributes to neuroinflammation. The role of glucose metabolism in glia to metabolically support neurons has been well established (Morrison et al., 2013; Pellerin and Magistretti, 1994; Chatterjee and Perrimon, 2021). In this study, we revealed an underappreciated role of glucose in the pigment glia to regulate its inflammatory responses.

Our study revealed that pigment glial cells can function as immune cells in the fly retina, which are activated in response to tau-induced lesions. Pigment glia ensheathes individual ommatidium, the minimum unit of the compound eye, to provide optical insulation that prevents extraneous light rays from inappropriately activating the photoreceptors (Tomlinson, 2012). It also accumulates lipid droplets in response to elevated oxidative stress in photoreceptor neurons (Liu et al., 2015), and lipid accumulation has been linked to microglial activation in mammalian microglia (Jung and Mook-Jung, 2020; Marschallinger et al., 2020). Our finding is consistent with the results from a comprehensive single-cell RNA sequence that AMP expression overlaps with a pigment glia marker “*santa-maria*” (Li et al., 2022; Yeung et al., 2022). Further characterization of pigment glia to identify the equivalent mammalian glial cell types will make the *Drosophila* retina an *in vivo* platform to study neuroinflammation and neuron-glia interaction.

We found that tau expression driven by *GMR-GAL4* causes swelling in the laminal cortex and degeneration of surface glia and cortex glia (Fig. 1), which *GMR-GAL4* is not expressed (Fig. S4). These non-cell autonomous effects may be mediated by AMPs, since GLUT expression in the pigment glia suppressed AMP expression and mitigated the laminal cortex swelling (Fig. 5). Although the primary functions of AMPs are to kill bacteria (Ganesan et al., 2011), recent reports suggest that AMPs affects diverse physiological events such as sleep and memory formation (Toda et al., 2019; Barajas-Azpeleta et al., 2018), suggesting AMPs can work as intercellular signaling molecules. In a fly model of traumatic brain injury, downregulation of the NF-κB signaling by a *Relish* mutation or loss of Mtk protects against detrimental effects (Swanson et al., 2020). Secreted AMPs from pigment cells in the tau-expressing retina may damage glial cells in the laminal cortex. Another possible mechanism underlying this non-cell-autonomous toxicity may be intercellular lipid transfer. During oxidative stress, neurons release lipids, which are internalized by glia and cause them to swell (Liu et al., 2015; Liu et al., 2017). Pigment cells primarily accept lipid droplets from photoreceptor neurons (Liu et al., 2015), while under neurodegenerative conditions, oxidative stress and lipid shuttling from neurons may exceed the capacity of the pigment cells, and overflowed lipids may harm distal glial cells. Enhancement of glucose uptake in pigment cells may enhance their ability to degrade lipid droplets from neurons and reduce the overflow of lipid droplets that affect other glia.

Enhanced glucose uptake in the pigment cells suppressed AMP expression dramatically; however, photoreceptor degeneration was only mildly rescued (Fig. 5). Tau affects many cellular components, such as the cytoskeleton, mitochondria, chromatin structures, lipids, and stress signaling, which can independently contribute to neurodegeneration (DuBoff et al., 2012; Frost et al., 2014; Fulga et al., 2007; Goodman et al., 2024; Papanikolopoulou et al., 2019; Voelzmann et al., 2016). Our results suggest that AMP secretion from pigment glia in the tau-expressing retina is one of the events induced in parallel in multiple cell types that collectively cause degeneration.

Changes in glial activity under disease conditions are often associated with alterations in the metabolic signatures, including glucose metabolism pathways (Edison, 2020; Jha and Morrison, 2018; Wang, 2019). Multiple signaling pathways that sense metabolic status are conserved across species and are also known to regulate inflammatory responses (Saito et al., 2019). For example, Liver kinase B1 (LKB1) is a key regulator of metabolism (Compton et al., 2023), and one of the downstream kinases of LKB1, AMP-activated protein kinase (AMPK), has been reported to promote anti-inflammatory responses (Compton et al., 2023) via negatively regulating the nuclear factor κB (NF-κB) signaling in microglia (Ramamurthy and Ronnett, 2006). Another critical energy regulator, Sirtuin 1 (SURT1), can suppress NF-κB signaling (Zhang et al., 2020). Many regulatory mechanisms of NF-κB signaling are conserved among species (Buchon et al., 2014), and these pathways can also regulate inflammatory responses in *Drosophila* (Hetru and Hoffmann, 2009). Multiple AMPs downstream of NF-κB signaling are activated upon tau-expression and suppressed upon GLUT3 co-expression. Further studies using metabolic and transcriptomic approaches will facilitate the mechanistic understanding of neuroinflammation.

## Materials and Methods

### Fly stocks and husbandry

The following fly stocks were obtained from the Bloomington *Drosophila* Stock Center (BDRC): *UAS-mCherry.NLS* (39434), *Repo-GAL4* (7415), *UAS-mCD8::ChRFP* (27391), *UAS-Luciferase* (35788), *UAS-Luciferase RNAi* (31603), *GMR-tau* P301L (51377), *UAS-CD8::GFP* (5137), *GMR-GAL4* (9146), *Rh1-GAL4* (8691), and *54C-GAL4* (27328). *UAS-Glut1^d05758^*, which is a P{XP} insertion containing UAS elements upstream of *Glut1*, was obtained from Harvard Medical School (http://flybase.org/reports/FBti0055936.htm, Exelixis *Drosophila* Collection) (Thibault et al., 2004). *UAS-drpr RNAi* (HMJ30231) and *UAS-NimC4 RNAi* (HMJ23355) were obtained from the Japanese National Institute of Genetics (NIG-FLY). *w^1118^* and *UAS-GLUT3* were a gift from Marie Thérèse Besson (Besson et al., 2015). *UAS-tau* (wildtype 0N4R) was a gift from Mel. B. Feany (Wittmann et al., 2001). Flies were reared in a standard medium containing 10% glucose, 0.7% agar, 9% cornmeal, 4% Brewer’s yeast, 0.3% propionic acid, and 0.1% n-butyl p-hydroxybenzoate [w/v]. Flies were maintained at 25°C under light-dark cycles of 12:12 hours. Experiments involving transgenic *Drosophila* were approved by the Tokyo Metropolitan University research ethical committee (#G5-15). Fly genotypes for each experiment are listed in Tables S1 and S2.

### Measurement of eye size

Images of fly eyes were captured with an Olympus digital microscope DSX110 or the Leica MZ16 stereomicroscope, and the area of the eye surface was measured using Fiji (NIH). Eyes from more than 11 flies were analyzed for each genotype.

### Immunostaining

Flies were collected at 3-5 days old, and brains were dissected in PBS, fixed in 4% PFA/PBS (Thermo Scientific) for 30 min at R.T. Brains were washed with PBST for 10 min three times. 4% normal donkey serum (abcam) was used as a blocking buffer for 30 min at R.T. Brains were stained with anti-elav (1:1000, DSHB Cat# Elav-9F8A9, RRID:AB_528217), anti-Repo (1:50, DSHB Cat# 8D12 anti-Repo, RRID:AB_528448), or anti-ubiquitin (FK2, 1:1000, StressMarq Biosciences Cat# SMC-214D-DY405, RRID:AB_2820835) and visualized with goat anti-rat antibody with Alexa Fluor 488 (1:1000, Thermo Fisher Scientific Cat# A-11006, RRID:AB_2534074)) or goat anti-mouse antibody with Alexa Fluor 488 (1:1000, (Thermo Fisher Scientific Cat# A-11001, RRID:AB_2534069). Nuclei were stained with DAPI (1:1000, abcam). Filamentous actin was stained with Alexa Flour 488 Phalloidin (1:1000, Invitrogen). Brains were mounted in VectaShield (Vector Laboratories) and imaged using Nikon spinning-disc confocal microscope.

### Western blotting

Flies were frozen with liquid nitrogen, and fifteen fly heads for each genotype were homogenized in the SDS-Tris-Glycine sample buffer. The same amount of the lysate was loaded to each lane of multiple 10% Tris-Glycine gels and transferred to PVDF membrane. The membranes were blocked with 5% skim milk in TBST and blotted with the antibodies described below, incubated with appropriate secondary antibodies, and detected using Immobilon Western Chemiluminescent HRP Substrate (Merck Millipore). The membranes were probed with anti-actin (sigma) as the loading control. Anti-tau pS202 (CP13) and anti-phospho-Ser396/404-tau antibody (PHF1) were a gift from Dr. Peter Davis (Albert Einstein College of Medicine, USA). Anti-tau antibody (T46, Thermo Fisher Scientific Cat# 13-6400, RRID:AB_2533025), Anti-pospho-Ser262-tau antibody (pSer262, abcam, Abcam Cat# ab92627, RRID:AB_10563129) anti-rhodopsin 1 antibody (4c5, DSHB Cat# 4c5, RRID:AB_528451), and anti-actin antibody (Sigma-Aldrich Cat# A2066, RRID:AB_476693) were purchased. The chemiluminescent signal was detected by Fusion FX (Vilber), and intensity was quantified using Fiji (NIH). Western blots were repeated a minimum of 3 times with different animals, and representative blots are shown. Flies used for western blotting were 5-7 days old after eclosion.

### Histological analysis

Fly heads at 3-5 days after eclosion were fixed in Bouin’s fixative for 48 hours at room temperature (R.T.) and incubated for 24 hours in a leaching buffer (50 mM Tris/150 mM NaCl). Samples were dehydrated with a series of ethanol baths (70%, 80%, 95%, and 100%) and xylene at R.T and embedded in paraffin. Serial sections (6 μm thickness) through the entire heads were stained with hematoxylin and eosin. Sections were mounted to glass slides, de-paraffined in xylene, and a series of ethanol baths (100%, 95%, and 80%) for 3min each. Slides were stained with 100% hematoxylin for 2 min and washed with water for 5 min. They were stained with 100% eosin for 30-45 seconds, washed with 95% ethanol and 100% ethanol for 6 min each, incubated with xylene for 10 min, dried, and mounted with a coverslip. Each section was examined by bright-field microscopy. Images of the sections that include the lamina were captured with the microscope BZ-X700 (Keyence) or the microscope Mica (Leica). The retinal degeneration or the vacuole area was analyzed using Fiji (NIH). The area of the laminal cortex was measured and normalized to the whole laminal area to calculate the laminal cortex area.

### Electron microscopy

Decapitated heads were cut in half vertically, then incubated in primary fixative solution (2.5% glutaraldehyde and 2% paraformaldehyde in 0.1 M sodium cacodylate buffer) at R.T. for 2 hours. After washing heads with 3% sucrose in 0.1 M sodium cacodylate buffer, fly heads were post-fixed for 1 hour in secondary fixation (1% osmium tetroxide in 0.1 M sodium cacodylate buffer) on ice. After washing with H2O, heads were dehydrated with ethanol and infiltrated with propylene oxide and Epon mixture (TAAB and Nissin EM) for 3 hours. After infiltration, specimens were embedded with an Epon mixture at 70°C for 2∼3 days. Some longitudinal sections cut out by glass knives were stained with toluidine blue to check the position. Thin sections (70 nm) of retinal sections were collected on copper grids. The sections were stained with 2% uranyl acetate in 70% ethanol and Reynolds’ lead citrate solution. Electron micrographs were obtained with a VELETA CCD Camera (Olympus Soft Imaging Solutions GMBH) mounted on a JEM-1010 electron microscope (Jeol Ltd.).

### RNA extraction and quantitative real-time PCR analysis

Heads from more than 25 flies were mechanically isolated, and total RNA was extracted using ISOGEN (NipponGene), followed by reverse transcription using the PrimeScript RT reagent kit (Takara). The resulting cDNA was used as a template for PCR with THUNDERBIRD SYBR qPCR mix (TOYOBO) on a Thermal Cycler Dice real-time system TP800 (Takara). Expression of genes of interest was standardized relative to *rp49* or *actin*. Relative expression values were determined by the ΔΔCT method (Livak and Schmittgen, 2001). Experiments were repeated three times, and a representative result was shown. Primers were designed using DRSC FlyPrimerBank (Harvard Medical School). Primer sequences are shown below.

*Attacin C* for 5’-CTGCACTGGACTACTCCCACATCA-3′

*Attacin C* rev 5’-CGATCCTGCGACTGCCAAAGATTG-3′

*Cecropin A1* for 5′-CATTGGACAATCGGAAGCTGGGTG-3′

*Cecropin A1* rev 5’-TAATCATCGTGGTCAACCTCGGGC-3′

*Diptericin B* for 5′-AGGATTCGATCTGAGCCTCAACGG-3′

*Diptericin B* rev 5’-TGAAGGTATACACTCCACCGGCTC-3′

*Drosomycin* for 5′-AGTACTTGTTCGCCCTCTTCGCTG-3′

*Drosomycin* rev 5’-CCTTGTATCTTCCGGACAGGCAGT-3′

*Metchnikowin* for 5′-CATCAATCAATTCCCGCCACCGAG-3′

*Metchnikowin* rev 5’-AAATGGGTCCCTGGTGACGATGAG-3′

*drpr* for 5’-GCAGATGCCTGAATAACTCCTC-3′

*drpr* rev 5’-TCCTTGCATTCCATGCCGTAG-3′

*NimC4* for 5’-GAACGAGACGATACGAGCCAC-3′

*NimC4* rev 5’-GGTGACTTGTTCCTCCTCTGA-3′

*Glut1* for 5’-TTACCGCGGAGCTCTTCTCC-3’

*Glut1* rev 5’-GCCATCCAGTTGACCAGCAC-3’

*rp49* for 5′-GCTAAGCTGTCGCACAAATG-3′

*rp49* rev 5′-GTTCGATCCGTAACCGATGT-3′

*actin 5C* for 5′-TGCACCGCAAGTGCTTCTAA-3′

*actin 5C* rev 5′-TGCTGCACTCCAAACTTCCA-3′

## Acknowledgments

We thank the Japanese National Institute of Genetics (NIG-FLY), Bloomington *Drosophila* Stock Center (BDSC), and Vienna *Drosophila* Resource Center (VDRC) for fly strains. We thank Dr. Shin-Ichi Hisanaga (Tokyo Metropolitan University) for critical comments and Drs. Taro Saito and Akiko Asada (Tokyo Metropolitan University) for technical support. We thank Drs. Mel B. Feany (Harvard University) for *UAS-tau* and Marie Thérèse Besson (Aix-Marseille Université) for *UAS-GLUT3*. We also thank Dr. Ismael Al-Ramahi (Baylor College of Medicine) for help with histological studies.

## Author contributions

Conceptualization: M.O. and K.A., Methodology: M.O. and K.A., Validation: M.O., Formal analysis: M.O., E. S. and K.A., Investigation: M.O. and S.N. Writing – original draft preparation: M.O. and K.A., Writing – review and editing: M.O., E.S., S.Y. and K.A., Visualization: M.O., Supervision: S.Y. and K.A, Project administration: S.Y. and K.A, Funding acquisition: M.O., S.Y. and K.A

## Data availability

All relevant data can be found within the article and its supplementary information.

---

**Figure S1. *UAS-Tau* alone does not reduce eye size**. The area of the eyes was similar between *GMR>Luciferase* and UAS-tau, and that of *GMR> Tau* was significantly less. Scale bar, 250 µm. Mean±SE, n=7-13, ***; p<0.001, n.s.; p>0.05, One-way ANOVA followed by Tuckey’s HSD multiple comparisons test.

**Figure S2. Knockdown of drpr or NimC4 in the tau-expressing retina does not affect eye size.** (A) Fly heads expressing *UAS-drpr* RNAi or *UAS-NimC4* RNAi driven by *GMR-GAL4* were subjected to qRT-PCR. Mean±SE, n=3, ***; p<0.001, Statistical significance was assessed with unpaired two-tailed t-test. (B) The eyes of flies co-expressing Tau and drpr RNAi or NimC4 RNAi. mCherry RNAi was used as a control. Scale bar, 250 µm. Mean±SE, n=15-17, n.s.; p>0.05, One-way ANOVA followed by Tuckey’s HSD multiple comparisons test.

**Figure S3. Overexpression of Glut1 with UAS-Glut1^d05758^ expression driven by *GMR-GAL4***. Heads of the flies with *UAS-Glut1^d05758^* expression driven by *GMR-GAL4* were subjected to qRT-PCR. Mean±SE, n=3, ***; p<0.001, Statistical significance was assessed with unpaired two-tailed t-test.

**Figure S4. Expression patterns of *GMR-GAL4* and *54C-GAL4***. *GMR-GAL4*- or *54C-GAL4*-expressing cells are labeled with *UAS-mCh::NLS* (magenta). Glial cells are immunostained by an anti-repo antibody (green). White dashed lines indicate the laminal cortex. Scale bar: 50 µm.

**Figure S5. The eyes with Tau expression driven by *54C-GAL4***. Tau expression was driven by 54C-GAL4. Luciferase was used as a control (*54C>Luciferase*). No significant difference was found in the area of the eye. Scale bar, 100 µm. Mean±SE, n=14-24, n.s.; p>0.05. Statistical significance was assessed with unpaired two-tailed t-test.

