## Supplementary material for "Glucose uptake in pigment glia suppresses tau-induced inflammation and photoreceptor degeneration in *Drosophila*": Supplemmentary

#### Supplementary information

**Figure S1-S5**

**Table S1-S2**

Figure S1

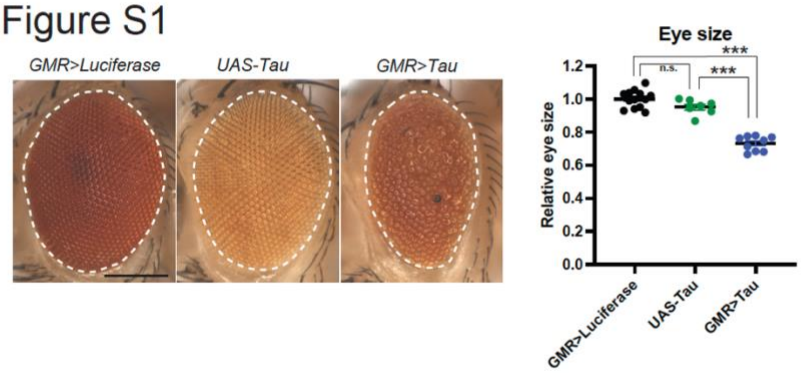


**Figure S1. *UAS-Tau* alone does not reduce eye size**. The area of the eyes were similar between *GMR>Luciferase* and UAS-tau, and that of *GMR>tau* was significantly less. Scale bar, 250 µm. Mean±SE, n=7-13, ***; $p<0.001$, n.s.; $p>0.05$, One-way ANOVA followed by Tuckey's HSD multiple comparisons test.

### Figure S2

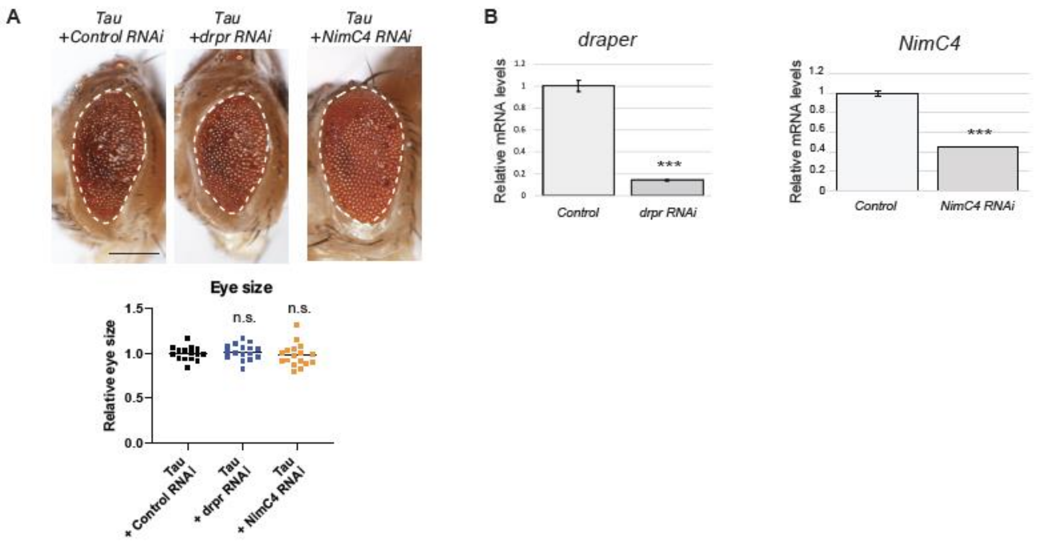


**Figure S2. Knockdown of Drpr or NimC4 in tau-expressing retina does not affect eye size.** (A) Fly heads expressing *UAS-drpr* RNAi or *UAS-NimC4* RNAi driven by *GMR-GAL4* were subjected to qRT-PCR. Mean±SE, n=3, ***; $p<0.001$, Statistical significance was assessed with unpaired two-tailed t-test. (B) The eyes of flies co-expressing tau and drpr RNAi or NimC4 RNAi. mCherry RNAi was used as a control. Scale bar, 250 µm. Mean±SE, n=15-17, n.s.; $p>0.05$, One-way ANOVA followed by Tuckey's HSD multiple comparisons test.

### Figure S3

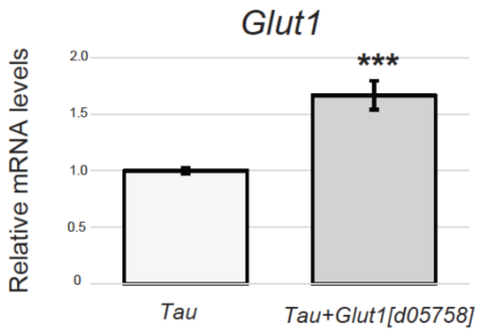


**Figure S3. Overexpression of GLUT1 with UAS-Glut1$^{d05758}$ expression driven by *GMR-GAL4*.** Heads of the flies with *UAS-Glut1$^{d05758}$* expression driven by *GMR-GAL4* were subjected to qRT-PCR. Mean±SE, n=3, ***; p<0.001, Statistical significance was assessed with unpaired two-tailed t-test.

### Figure S4

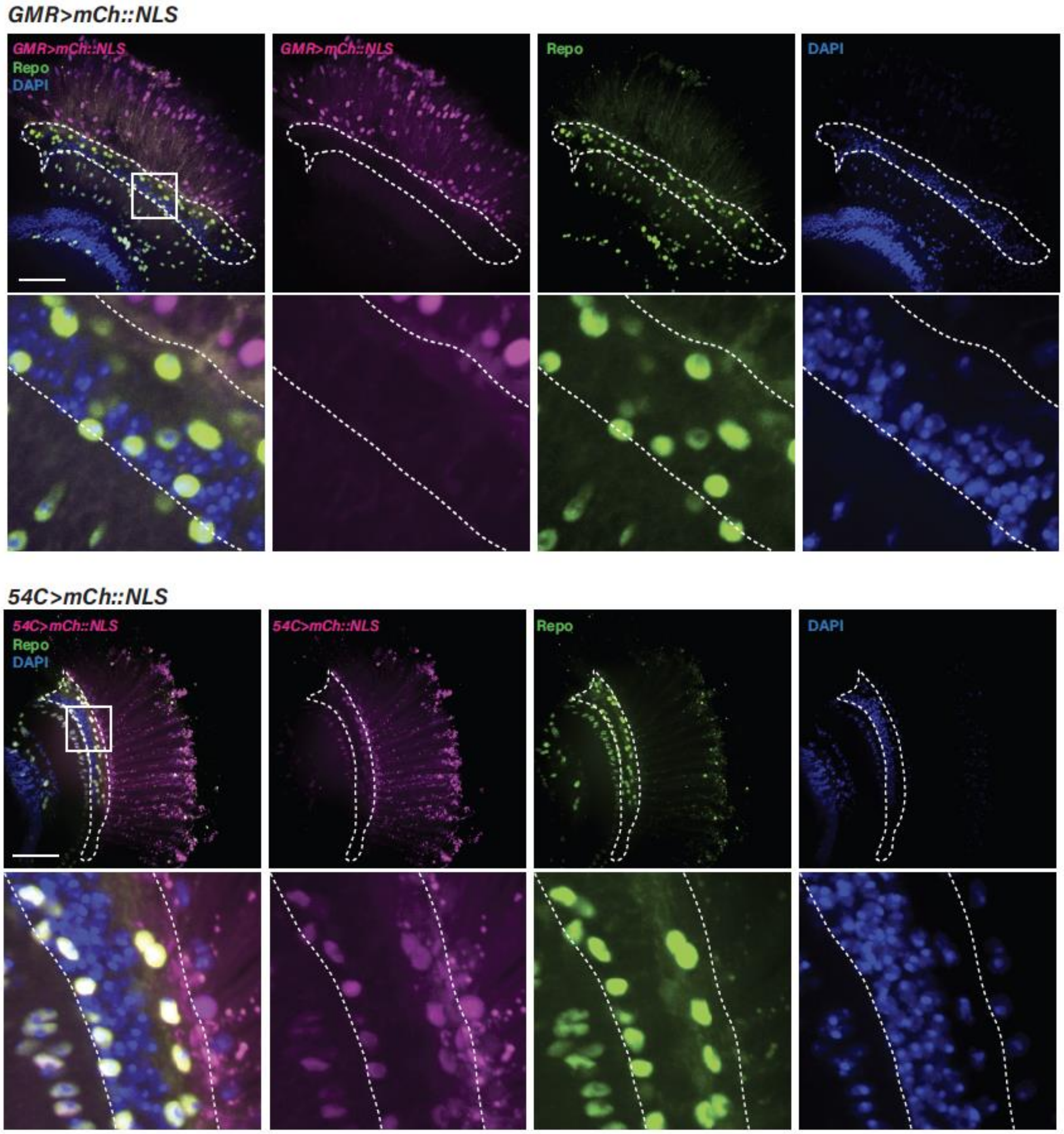


**Figure S4. Expression patterns of *GMR-GAL4* and *54C-GAL4***. *GMR-GAL4*- or *54C-GAL4*-expressing cells are labeled with expression of *UAS-mCh::NLS* (magenta). Glial cells are immunostained by anti-repo antibody (green). White dashed lines indicate the laminal cortex. Scale bar; 50 µm.

Figure S5

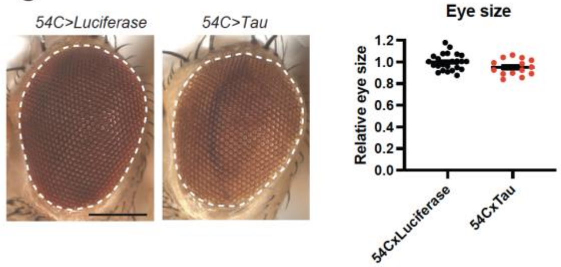


**Figure S5. The eyes with Tau expression driven by *54C-GAL4***. Tau expression was driven by 54C-GAL4. Luciferase was used as a control (*54C>Luciferase*). No significant difference was found in the area of the eye. Scale bar, 250 μm. Mean±SE, n=14-24, n.s.; p>0.05, Statistical significance was assessed with unpaired two-tailed t-test.

**Table S1. The genotypes of flies used in each experiment related to Figure 1-5**

| Figure | Name | *Genotype* |
|---|---|---|
| Figure 1A-C, 1E-F and G, left | Control | *$w^{1118}$; GMR-GAL4/+* |
| | Tau | *$w^{1118}$; GMR-GAL4/+; UAS-tau/+* |
| Figure 1D | Control | *$w^{1118}$;; Repo-GAL4/UAS-mCh::NLS* |
| | Tau | *GMR-Tau/+;; Repo-GAL4/UAS-mCh::NLS* |
| Figure G, right | Tau | *GMR-Tau/+* |
| Figure 2A | Tau+Control RNAi | *$w^{1118}$; GMR-GAL4/+; UAS-tau/UAS-Luciferase RNAi* |
| | Tau+drpr RNAi | *$w^{1118}$; GMR-GAL4/UAS-drpr RNAi; UAS-tau/+* |
| | Tau+NimC4 RNAi | *$w^{1118}$; GMR-GAL4/UAS-NimC4 RNAi; UAS-tau/+* |
| Figure 2B | Control | *$w^{1118}$; GMR-GAL4/+* |
| | Tau | *$w^{1118}$; GMR-GAL4/+; UAS-tau/+* |
| Figure 3A | Control | *$w^{1118}$; GMR-GAL4/+* |
| | Tau | *$w^{1118}$; GMR-GAL4/+; UAS-tau/+* |
| | Tau+GLUT3 | *$w^{1118}$; GMR-GAL4/UAS-GLUT3; UAS-tau/+* |
| | Tau+CD8::RFP | *$w^{1118}$; GMR-GAL4/UAS-mCD8::ChRFP; UAS-tau/+* |
| | GLUT3 | *$w^{1118}$; GMR-GAL4/UAS-GLUT3* |
| Figure 3B-F | Control | *$w^{1118}$; GMR-GAL4/+* |
| | Tau | *$w^{1118}$; GMR-GAL4/+; UAS-tau/+* |
| | Tau+GLUT3 | *$w^{1118}$; GMR-GAL4/UAS-GLUT3; UAS-tau/+* |
| Figure 3G-I | Control | *$w^{1118}$; GMR-GAL4/UAS-mCD8::ChRFP* |
| | Tau | *$w^{1118}$; GMR-GAL4/ UAS-mCD8::ChRFP; UAS-tau/+* |
| | Tau+Glut1$^{d05758}$ | *$w^{1118}$; GMR-GAL4/+; UAS-tau/ UAS- Glut1$^{d05758}$* |
| | Glut1$^{d05758}$ | *$w^{1118}$; GMR-GAL4/+; UAS- Glut1$^{d05758}$/+* |
| Figure 4 | Tau | *$w^{1118}$; GMR-GAL4/+; UAS-tau/+* |
| | Tau+GLUT3 | *$w^{1118}$; GMR-GAL4/UAS-GLUT3; UAS-tau/+* |
| Figure 5A-D | 54C>CD8::GFP | *$w^{1118}$; 54C-GAL4/ UAS-mCD8::GFP* |
| | GMR-Tau +54C>CD8::GFP | *GMR-Tau/+; 54C-GAL4/UAS-mCD8::GFP* |
| | GMR-Tau | *GMR-Tau/+; 54C-GAL4/UAS-GLUT3* |

| | | |
|---|---|---|
| | +54C>GLUT3 | |
| | 54C>GLUT3 | *$w^{1118}$; 54C-GAL4/UAS-GLUT3* |
| Figure 5E-G | Rh1>CD8::GFP | *$w^{1118}$; UAS-mCD8::GFP /+; Rh1-GAL4/+* |
| | GMR-Tau +Rh1>CD8::GFP | *GMR-Tau/+; UAS-mCD8::GFP/+; Rh1-GAL4/+* |
| | GMR-Tau +Rh1>GLUT3 | *GMR-Tau/+; UAS-GLUT3/+; Rh1-GAL4/+* |
| | Rh1>GLUT3 | *$w^{1118}$; UAS-GLUT3/+; Rh1-GAL4/+* |

**Table S2. The genotypes of flies used in each experiment related to Supplemental Figure 1-5**

| Figure | Name | *Genotype* |
|---|---|---|
| Supplemental Figure 1 | GMR>Luciferase | *$w^{1118}$; GMR-GAL4/+; UAS-Luciferase/+* |
| | UAS-Tau | *$w^{1118}$;; UAS-Tau/+* |
| | GMR>Tau | *$w^{1118}$; GMR-GAL4/+; UAS-Tau/+* |
| Supplemental Figure 2 | Control | *$w^{1118}$; GMR-GAL4/+; UAS-Luciferase RNAi/+* |
| | drpr RNAi | *$w^{1118}$; GMR-GAL4/UAS-drpr RNAi* |
| | NimC4 RNAi | *$w^{1118}$; GMR-GAL4/UAS-NimC4 RNAi* |
| Supplemental Figure 3 | Tau | *Elav-GAL4/+; UAS-mCD8::GFP/+; UAS-tau/+* |
| | Tau+Glut1 | *Elav-GAL4/+;; UAS-tau/ UAS- $Glut1^{d05758}$* |
| Supplemental Figure 4 | GMR>mCh::NLS | *$w^{1118}$; GMR-GAL4/+; UAS-mCh::NLS/+* |
| | 54C>mCh::NLS | *$w^{1118}$; GMR-GAL4/54C-GAL4* |
| Supplemental Figure 5 | 54C>Luciferase | *$w^{1118}$; 54C-GAL4/+; UAS-Luciferase/+* |
| | 54C>Tau | *$w^{1118}$; 54C-GAL4/+; UAS-Tau/+* |
